# Directed functional connectivity using dynamic graphical models

**DOI:** 10.1101/198887

**Authors:** Simon Schwab, Ruth Harbord, Valerio Zerbi, Lloyd Elliott, Soroosh Afyouni, Jim Q. Smith, Mark W. Woolrich, Stephen M. Smith, Thomas E. Nichols

## Abstract

There are a growing number of neuroimaging methods that model spatio-temporal patterns of brain activity to allow more meaningful characterizations of brain networks. This paper proposes dynamic graphical models (DGMs) for dynamic, directed functional connectivity. DGMs are a multivariate graphical model with time-varying coefficients that describe instantaneous directed relationships between nodes. A further benefit of DGMs is that networks may contain loops and that large networks can be estimated. We use network simulations, human resting-state fMRI (N = 500) to investigate the validity and reliability of the estimated networks. We simulate systematic lags of the hemodynamic response at different brain regions to investigate how these lags potentially bias directionality estimates. In the presence of such lag confounds (0.4–0.8 seconds offset between connected nodes), our method has a sensitivity of 72%–77% to detect the true direction. Stronger lag confounds have reduced sensitivity, but do not increase false positives (i.e., directionality estimates of the opposite direction). In human resting-state fMRI, we find the DMN has consistent influence on the cerebellar, the limbic and the auditory/temporal network, as well a consistent reciprocal relationship between the visual medial and visual lateral network. Finally, we apply the method in a small mouse fMRI sample and discover a highly plausible relationship between areas in the hippocampus feeding into the cingulate cortex. We provide a computationally efficient implementation of DGM as a free software package for R.

## Introduction

Human behavior is underpinned by brain circuits and there is evidence of abnormal connectivity in brain networks related to the psychopathology of mental disorders (Bassett and Bullmore, 2009; Buckner et al., 2008; Menon, 2011), for example, in schizophrenia (Zhou et al., 2016), major depression (Cheng et al., 2016; Guo et al., 2016), dementia (Chase, 2014), or anxiety disorders (Peterson et al., 2014). New treatment strategies in the the future may benefit from new computational method developments that lead to an improved understanding of altered functional networks of the brain (Friston et al., 2014b; Huys et al., 2016; Stephan and Mathys, 2014). However, a majority of studies in functional connectivity (FC) are based on correlations (full, or partial) between fMRI time series, and even though they are fast to compute and can handle large networks, these have some major limitations (Smith et al., 2013b).

First, FC in terms of correlation between time series reflects static, stationary relationships with fixed connection strengths across time. However, coupling between brain systems is not constant over time—there is evidence that brain connectivity is better described in terms of time-varying connectivity (Allen et al., 2014; Liu and Duyn, 2013; Shine and Poldrack, 2017; Smith et al., 2012a). Emerging evidence suggests that dynamic functional connectivity may reflect changes in the underlying effective connectivity, where changes in effective connectivity cause—or are caused by—crucial aspects of cognition and behaviour (Allen et al., 2014; Braun et al., 2015; Hutchison et al., 2013a; Smith et al., 2012b; Zalesky et al., 2014). The study of these processes can be highly relevant in psychiatry (Damaraju et al., 2014; Kaiser et al., 2016; Zhang et al., 2016). Such dynamic coupling can occur in different behavioral contexts and during cognitive control (Cocchi et al., 2013; Gonzalez-Castillo et al., 2015; Shine et al., 2016). In particular, intermodular connections seem to be dynamic, linking different systems of the brain (Zalesky et al., 2014). Hence, it is more plausible that the brain exhibits dynamic communication (Andrews-Hanna et al., 2014; Betzel et al., 2016; Bressler and Kelso, 2001; Fox et al., 2005; Hutchison et al., 2013b). Only recently have studies begun to look closer at fluctuating dynamic connectivity, but proper validation is important (Laumann et al., 2017; Leonardi and Van De Ville, 2015; Lindquist et al., 2014; Ryali et al., 2011). Consequently, dynamic methods will potentially have greater impact than static methods in the future (Bassett and Gazzaniga, 2011; Calhoun et al., 2014; Preti et al., 2017; Vidaurre et al., 2017).

Second, FC in terms of a correlation between nodes reflects undirected relationships between them. A major challenge is to estimate the direction of information flow in FC (Ramsey et al., 2010). Notably, the rate of measurement for fMRI is much slower than the neural time lag, and the hemodynamic response (HR) is subject to variation of the temporal delay. Even though most variation is between subjects, there is also variation between brain regions (Handwerker et al., 2004) which can be problematic As a result, lag-based methods, such as Granger causality, can be confounded by the variability of the HR in different brain areas (Smith et al., 2013b, 2011). For example, if a node A is influencing a node B, but node A has a slower HR compared to B, lag-based methods may estimate the incorrect direction. An interesting issue here is the nature of the lag induced by different haemodynamic response functions in different nodes. Simply changing the parameters or shape of the haemodynamic response function does not necessarily invalidate Granger causality (Barnett and Seth, 2017). However, the introduction of explicit conduction delays violates the temporal precedence assumptions that underlie Granger causality.

Several methods have been proposed to infer causal relationships between nodes, for example exploiting non-Gaussian features of the BOLD signal (Hyvärinen and Smith, 2013; Ramsey et al., 2014) to determine the edge direction between a node pair. The IMaGES (Ramsey et al., 2011) method can successfully determine directionality on the group level with concatenated time series across subjects (multi-subject approach), and the GIMME algorithm recovers individual network structure after estimating the group structure (Gates and Molenaar, 2012). A very popular method to estimate directed networks is dynamic causal modeling (DCM; Friston et al., 2003). DCM is primarily designed to make inferences about how experimental conditions change connectivity (Stephan et al., 2010). When used to fit resting state time series, DCM is usually deployed over sliding windows to estimate fluctuations in effective connectivity (Cooray et al., 2016). Often, only small networks are investigated in DCM studies, as investigators have specific hypotheses on a small number of connections. However, new variants of DCM can be applied to large graphs (rDCM; Frässle et al., 2017; Seghier and Friston, 2013). For example, spectral DCM is suitable to study networks with dozens of nodes and is designed for resting-state experiments (Friston et al., 2014a; Razi et al., 2017, 2015).

In the present study, we use dynamic graphical models (DGM) to estimate dynamic, directed functional connectivity. There is overwhelming evidence that brain areas are not acyclic due to reciprocal polysynaptic connections (Friston, 2011; Markov et al., 2014). Therefore, unlike our related work on “multiregression dynamic models” (MDMs; Costa et al., 2015a), we do not constrain the networks to be acyclic. Without the acyclic constraint, we do not have a single statistical model for all nodes, but a heuristic approach, comprised of a set of dynamic linear models, one per node. We refer to our approach as the Dynamic Graphical Model (DGM). A DGM is a dynamic network with time-varying connection strengths that can accommodate larger directed networks. Here, “dynamic” refers to time-varying coupling strengths modeled (as detailed below) by a random walk model of the underlying coupling. In distinction to DCM, we do not consider any time varying experimental conditions or designed input. While this dynamic model is stationary, e.g. random walk smoothness parameters are constant in time, it can explain time-varying correlations in observed fMRI time series. Further, DGM have added interpretability as a regression model: for each child node, its time series are regressed on the time series from one or more other parent nodes; the directed relationship corresponds to the information flowing from the parent to the child node. Crucially, DGM use instantaneous interactions between the brain regions rather than lagged relationships to estimate directionality.

A major challenge of DGM is the number of possible graphs which increase exponentially as a function of the number of nodes. Searching such a large solution space is not feasible, even with modern computational resources. For example, for *n* = 15 regions/nodes there 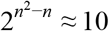 possible directed graphs^1^. However, the DGM model evidence factorizes by node (see below), and hence models can be optimised for each node independently, and thus only 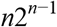 model evaluations (e.g. 245,760 for *n* = 15) are required. Even so, networks larger than 25 nodes are computationally intensive and could require several months to compute (Figure 1C), but, network discovery (stepwise model selection) are promising to drastically speed up the computational time in the future (Harbord et al., 2016). Also, we show that post-hoc pruning of the networks can further improve directionality estimates; such optimizations are an efficient way to explore model spaces and improve the computational burdens. (Friston et al., 2016; Seghier and Friston, 2013).

**Figure 1.**
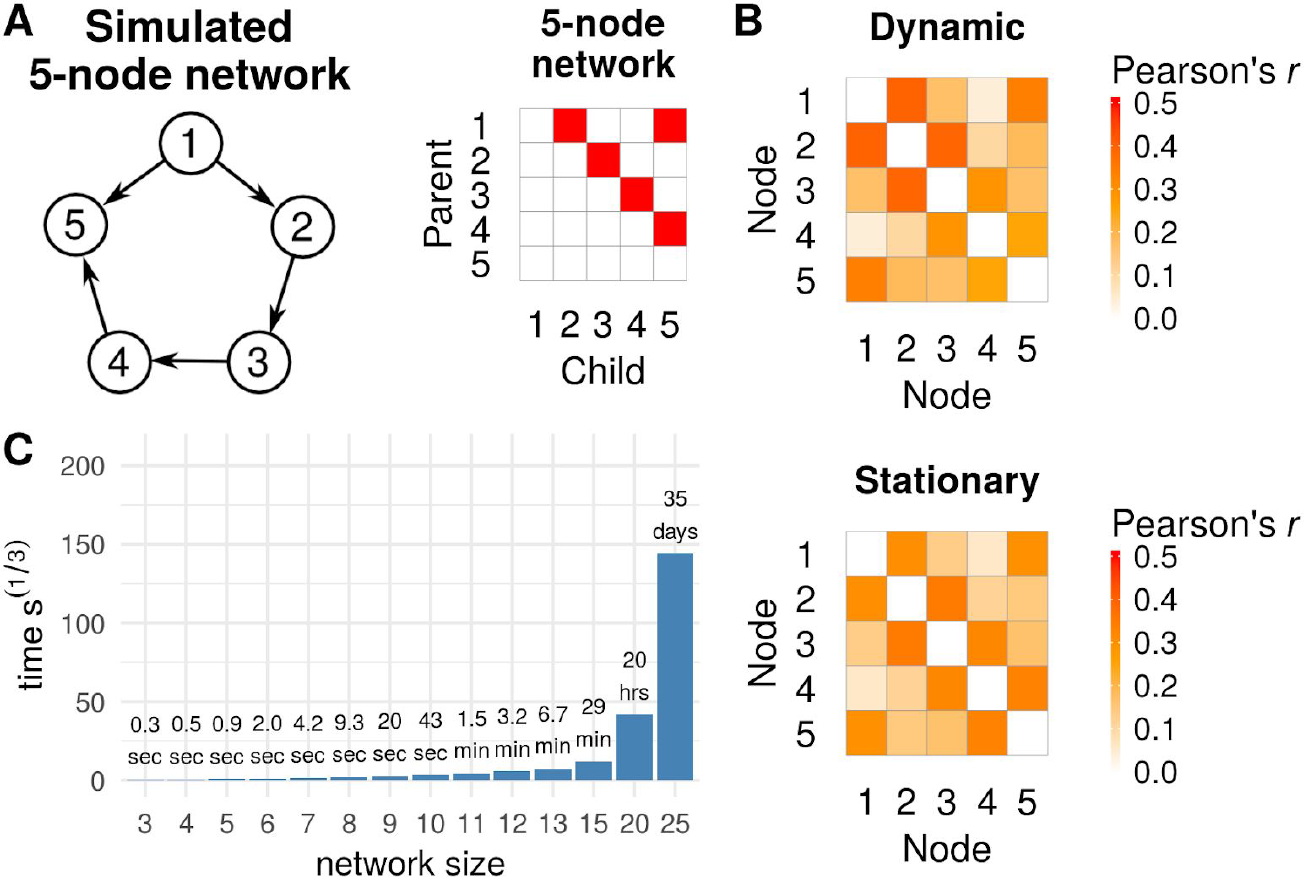
(A) True 5-node network and corresponding adjacency matrix. **(B)** Mean Pearson correlations across 50 simulations for the dynamic (Sim22, top) and stationary network simulations (Sim1, bottom). **(C)** Computational time estimates for a single node of a given network size for a time series of length 1,200 using a Intel Xeon CPU (E5-2630 v2) with 2.6 GHz.

The aim of this study is (1) to investigate the accuracy of DGM for recovering the true network structures in simulation based data, (2) to look at the reliability and consistency DGM networks in large samples of real fMRI data, (3) to give a worked example with mouse data and evaluate whether the network structures are plausible and relate to existing literature, and (4) to provide a fast implementation of DGM as an “R” package. Additionally, we include a post-hoc optimization that improves directionality estimates. For the synthetic data, we use generated data from network simulations that have similar properties as real BOLD data (Smith et al., 2011), as we aim to compare network estimates to a ground truth in terms of sensitivity and specificity. We further compare DGM’s performance to other directed functional connectivity methods. In subsequent work, we will extend this comparative evaluation to effective connectivity and DCM. At present, we are concerned with establishing the face validity of DGM; namely, its performance in terms of identifying true patterns of connectivity in relation to alternative functional connectivity methods. In subsequent work, we will address DGM’s ability to discriminate among different models of connectivity using Bayesian model comparison (although, a form of Bayesian model comparison is used implicitly to optimise DGM in the procedure described below). We hypothesize that DGM has better detection performance specifically with non-stationary data as the method can model the dynamic relationships between nodes. We also change the simulated node’s hemodynamic lags and hypothesize that the DGM, which is based on instantaneous dependencies between nodes, is not affected by variation in the haemodynamic response which can confound and reverse directionality estimates, for example as in purely lag-based methods. Having established the validity of DGM, we estimate dynamic directed relationships of resting-state networks (RSNs) in large samples from the Human Connectome Project to look at the reproducibility of the estimated networks. Finally, we investigate a hippocampal and a somatosensory network in mouse resting-state fMRI, where we have a clear expectation of the directionality based on findings from viral tracing studies in the mouse brain.

## Materials and methods

### Network simulations

We created various synthetic fMRI datasets based on network simulations of a 5-node (*N* = 50 datasets) and a 8-node network (*N* = 20), see Table 1. The simulated data follow the same approach as Smith et al. (2011), using a forward model (Friston et al., 2003) that links neural populations to a nonlinear balloon model (Buxton et al., 1998); for full details see Smith et al. (2011). In brief, each node has a binary external input consisting of a Poisson process with an added noise of 1/20 of the neural amplitudes. Mean duration of the simulated firing was 2.7 s (on) and 10 s (off); with a range between 770 ms and 10.1 s. The network also included self-connections to model within-node temporal decay; this was set to approximately 50 ms for the 5-node network, and 500 ms for the 8-node network. These specifications of neural activity is rather ad hoc, but these capture representative settings suitable for our evaluations. Each node’s neural signal was transformed into a simulated BOLD signal using the balloon model resulting in data with approximately 4% signal change, a signal that corresponds to typical fMRI data. These simulations had a hemodynamic lag variability of 0.5 s, reflecting observed variation of the HR from different brain regions. The first two simulated 5-node networks (Figure 1A) were “Sim1” and “Sim22” from Smith et al. (2011).^2^ Sim1 is a stationary network, and Sim22 an non-stationary (dynamic) network with time-dependent connection strengths. Both network’s time series were sampled to create 10 minute time series with a TR of 3 s. In Sim22 the connection strengths are modulated over time by additional random processes, such that the connection strength between two nodes is randomly set to zero for random intervals of mean duration of 30 seconds. For dynamic time series, correlation coefficients between truly connected nodes ranged from 0.25–0.40 across the 5 nodes (mean 0.33), and for stationary time series from 0.30–0.35 (mean 0.32), see Figure 1B.

**Table 1:** Specifications of the simulated BOLD-fMRI data. Each of the simulations contain a simulated 5-node networks for 50 simulated subjects.

| Simulation<br>Name | Number<br>of nodes | Neural<br>lag | Dynamic<br>network<br>strengths | Duration | TR | Noise | HRF lag variability<br>(s) |
| --- | --- | --- | --- | --- | --- | --- | --- |
| Sim1 | 5 | 50 ms | No | 10 min. | 3 s | 1.0% | $\pm 0.5^1$ |
| Sim22 | 5 | 50 ms | Yes | 10 min. | 3 s | 0.1% | $\pm 0.5^1$ |
| Offset < 0.4 s | 5 | 50 ms | Yes | 10 min. | 2 s | 0.1% | $\pm 0.3^1$ |
| Offset 0.4 s | 5 | 50 ms | Yes | 10 min. | 2 s | 0.1% | $+0.21/-0.20^2$ |
| Offset 0.8 s | 5 | 50 ms | Yes | 10 min. | 2 s | 0.1% | $+0.48/-0.35^2$ |
| Offset 1.1 s | 5 | 50 ms | Yes | 10 min. | 2 s | 0.1% | $+0.69/-0.46^2$ |
| Offset 1.4 s | 5 | 50 ms | Yes | 10 min. | 2 s | 0.1% | $+0.85/-0.54^2$ |
| Offset 1.7 s | 5 | 50 ms | Yes | 10 min. | 2 s | 0.1% | $+1.04/-0.58^2$ |
| Offset 1.9 s | 5 | 50 ms | Yes | 10 min. | 2 s | 0.1% | $+1.24/-0.59^2$ |
| 60 min. | 5 | 50 ms | Yes | 60 min. | 2 s | 0.1% | $\pm 0.3^1$ |
| 8-nodes ST | 8 | 500 ms | No | 10 min. | 2 s | 0.1% | $\pm 0.5^1$ |
| 8-nodes NS | 8 | 500 ms | Yes | 10 min. | 2 s | 0.1% | $\pm 0.5^1$ |

<sup>1</sup> standard deviation of HRF peak.
<sup>2</sup> average increase (node 1 and 4) / average decrease (node 4 and 5) of the hemodynamic delay.

We created seven additional simulations with systematic offsets between the nodes using the “NetSim” framework of Smith et al. (2011). With “offset”, we refer to the difference between two nodes with respect to their hemodynamic delays. For example, if we increase the HR delay at node 1 by +0.5 s and decrease the delay at node 2 by -0.2 s, the resulting total offset introduced between the two nodes is 0.7 s. The HR delay is the increased/decreased lag compared to the canonical HRF at a node, which is usually around 4–6 s. While Smith et al. (2011) randomly changed the delay of each node with a standard deviation of ±0.5 s, we *systematically* altered the HRF delay at four of the five nodes to create a worst-case, “slow parent, fast child” scenario for a lag-based method (Figure 2A). To achieve this, we changed the shape of the hemodynamic response function: we changed the transit time of blood τ in the balloon model, i.e., the average time blood takes to traverse the venous compartment (Buxton et al., 1998; Stephan et al., 2007). Values for τ were sampled from a normal distributions with means provided in the Supplementary Table S2 and a variance of 0.05. For node 1 and 4, we increased τ, and for node 2 and 5 we decreased τ. We confirmed this by a simulation using a single block of neural spiking with a duration of 2.5 s and measured the time to peak in the simulated HR at each node as the signal moves through the network; see Figure 2C and Supplementary Table S3. As a result our interventions increased node 1’s delay, and decreased node 2’s delay; the same intervention was performed for node 4 and node 5, and node 3 remained unaffected (Figure 2AC). Therefore, nodes 2 and 5 had their HRF peaks occurring before their respective parent nodes 1 and 4. In our simulations that included these offsets, node 1 and 4 had increased delays by 0.21–1.24 s compared to the canonical HRF shape, and node 2 and 5 had decreased delays by 0.20–0.59 s. These seven additional simulated datasets (Table 1) primarily differed regarding the strength of intervention in terms of total systematic offset between connected nodes, with the first dataset having no systematic intervention with an offset of < 0.4 s, and the other simulations having systematic offsets of 0.4 s, 0.8 s, 1.1 s, 1.4 s, 1.7 s, and 1.9 s. Additional descriptive statistics of these datasets are reported in Supplementary Tables S3–S5. The simulated nodes’ correlations were comparable to the Sim22 dataset (Figure 2B), and with increased intervention strength, the correlations decreased, as expected. Time series had a mean correlation of *r* = 0.32 across the five node-pairs and interventions. We also created a long 60 min. simulation of dynamic time series as session duration improved sensitivity in various network modeling methods (Smith et al., 2011).

**Figure 2.**
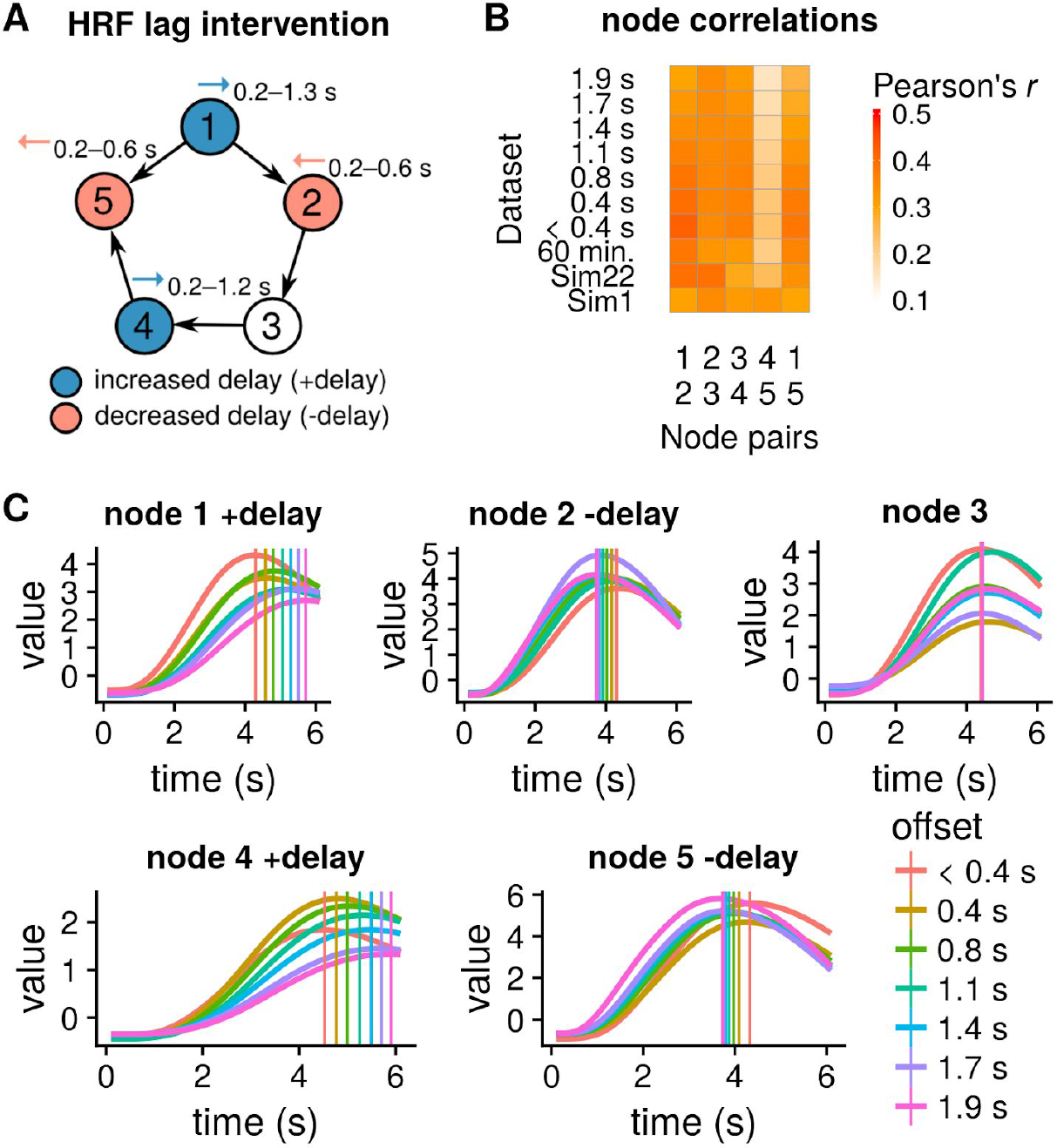
(A) Systematic interventions at four of the five nodes with either increased (node 1 and 4) or decreased HRF delay (node 2 and 5). We created seven simulations with a total offset between two nodes from 0.4–1.9 s **(B)** Pairwise node correlations in the different simulations. **(C)** Signal output at the 5 nodes showing the BOLD amplitude for the seven interventions where different DCM forward model parameters were used to shift peaks of the HR response, with colors from red (< 0.4 s offset) to pink (1.9 s; maximal intervention). Vertical lines indicate average time to peak after stimulus onset for each intervention type.

Finally, we extended the 5-node network and created a stationary and nonstationary 8-node network simulation with two reciprocal connections, and a neural lag of 500 s (Figure 7A). We aimed to investigate whether DGM can successfully detect the bidirectional relationships. The neural lag was changed from 50 ms to 500 ms as this more reasonably reflects neural activity by populations of neurons with slower fluctuations.

### fMRI data

For real fMRI data, we used resting state network (RSN) time series of 500 participants (57% female; age distribution of 42%, 36%, 21% and 1% in ranges of 26–30, 31–35, 22–25, and 36+ years) provided by the Human Connectome Project (Smith et al., 2013a; Van Essen et al., 2013). RSN time-series were from the HCP 900 d=25 “PTN” data (Parcellation, Time-series + Netmats), a 25-dimension ICA decomposition of the 15 minutes of resting state data which was acquired with a TR of 0.72 s. From the 25 spatial components we selected ten RSNs which had the highest spatial correlation with the ten RSNs reported by Smith et al. (2009), see Supplementary Table S1. Note that we are treating a RSN as a node in our analyses, i.e., we are summarising the activity of a distributed mode (i.e., resting state network) with a single time series and then using the DGM to understand how ’networks’ are coupled to ’networks’. The reason we can do this is that most resting state networks comprise spatially compact regions that can be treated as a small collection of nodes. For mouse rs-fMRI data, we analyzed two networks from a previously published dataset of 16 mice, the first is an amygdala-hippocampal-entorhinal network (8 nodes), the second a cortico-striatal-pallidum network with 6 nodes (Sethi et al., 2017; Zerbi et al., 2015).

### Dynamic Graphical Models

Dynamic Graphical Models (DGM) are graphical models with directed relationships and time-varying connectivity weights. The connectivity weights are the regression coefficients that reflect the effect of a set of parent nodes as covariates on a child node; see Figure 3. DGM are closely related to the Multiregression Dynamic Model (Costa et al., 2015b; MDM; Queen and Smith, 1993), except that a DGM need not be a directed acyclic graph (DAG). A DGM is a set of dynamic linear models (DLMs), state space models that are linear and Gaussian (Petris et al., 2009a; West and Harrison, 1997). In a DLM, the time series of a given child node are regressed on the time series from one or more other parent nodes using dynamic regression. The directed relationship corresponds to information flowing from the parent to the child. A DGM is notable for its computational tractability: conditional on an easily estimated ‘discount factor’, the model is fully conjugate and the model evidence available in closed form.

**Figure 3.**
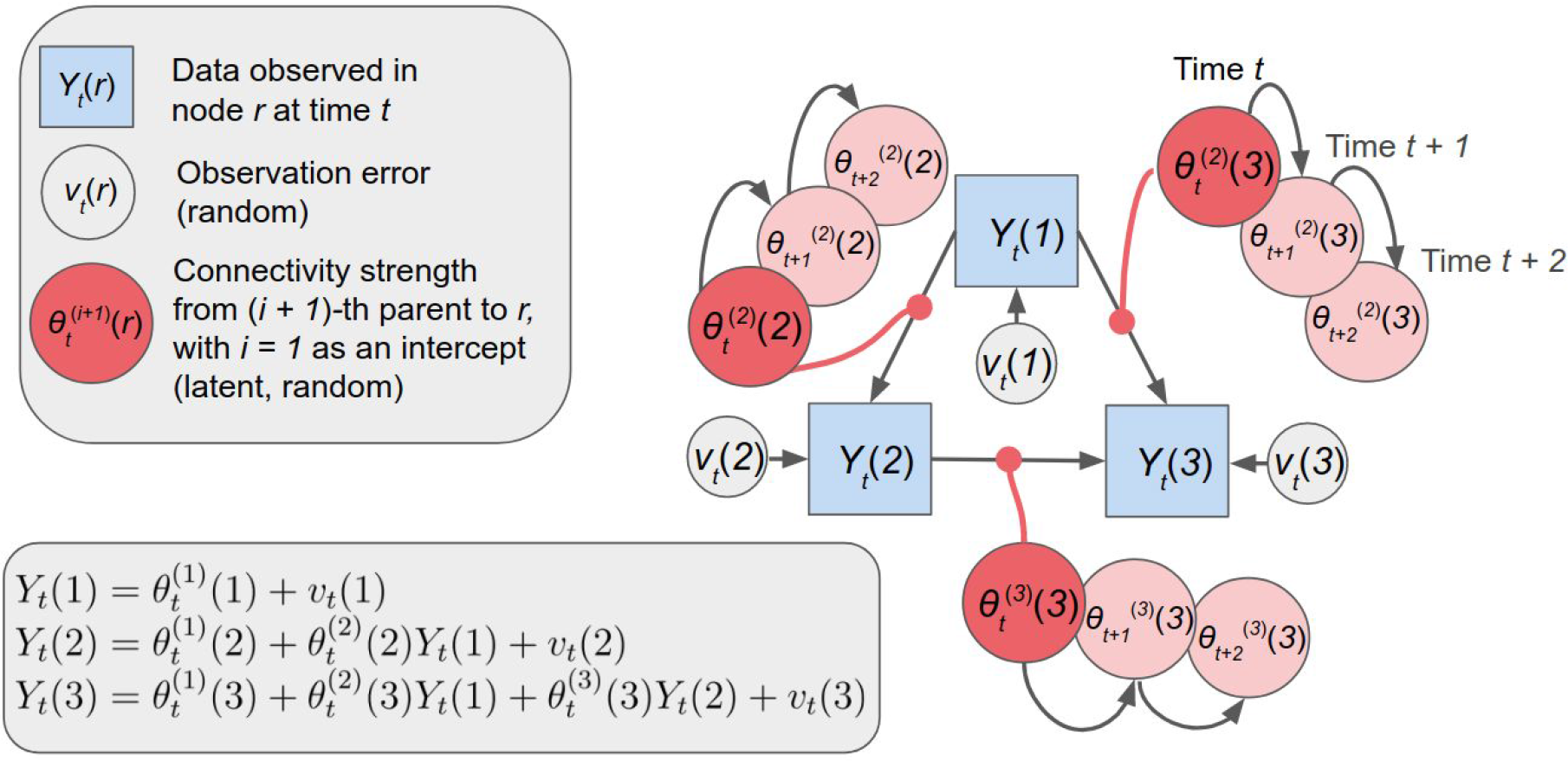
Dynamic Graphical Model (DGM) is a graphical model with directed relationships. θ*_t_*^(^*^i^*^)^(*r*) are the time-varying connectivity strength of parent region *i* + 1 on child node r (*i* = 1 is the intercept), or in other words, the varying regression coefficients from a dynamic regression. In this 3-node network, the model equations for the three nodes are written in the lower left box, showing the DGM to be a collection of dynamic linear models (DLM).

We define a DGM as a collection of multiple regression DLMs described by the following set of equations (Harrison and Stevens, 1976; West and Harrison, 1997; p. 109):

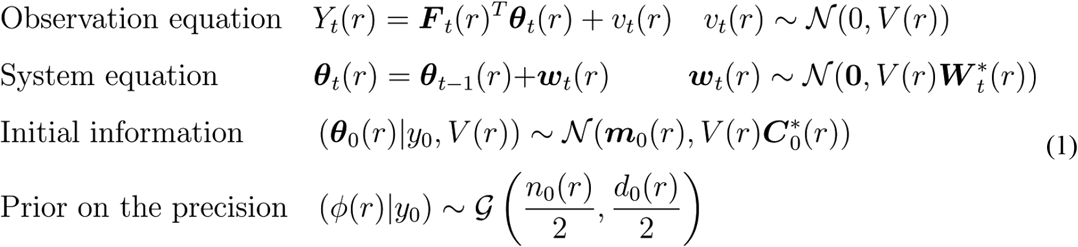

For some child region *r, Y_t_*(*r*) is the observed value at time *t*, (*r* = 1,…,*n*, *t* = 1,…,*T*). Let *Pa*(*r*) denote the parents of *r* and |Pa(*r*)| be the number of parents. Then ***θ****_t_*(*r*) is an unobservable system vector with length |*Pa*(*r*)| *+ 1*. The covariate vector ***F****_t_*(*r*) has the same dimension as ***θ****_t_*(*r*) and contains the observations from the parent nodes *Pa*(*r*) at time *t*; the first element of ***F****_t_*(*r*) is 1 to provide an intercept. The observation error *v_t_*(*r*) and the system error ***w****_t_*(*r*) are normally-distributed with variances *V*(*r*) and 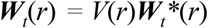 respectively. The error vectors are assumed to be mutually independent and independent over time and nodes. The observation variance *V*(*r*) is unknown and assumed to be constant so that we may place a gamma-distributed prior on the precision ɸ(*r*) *= V*(*r*)*^-1^* and express all of the variances in the DLM in terms of this constant, unitless term (West and Harrison, 1997). Finally, the initial system vector ***θ***_0_(*r*), conditioned on *y*_0_, is the available information at time *t* = 0, and represents prior knowledge of the regression coefficients before observing any data; ***θ****_0_*(*r*) follows a normal distribution with mean ***m***_0_(*r*) and variance 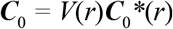.

The system variance is re-parameterized in terms of the posterior variance 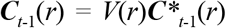 of the state variables ***θ****_t_*(*r*) at time *t* - 1

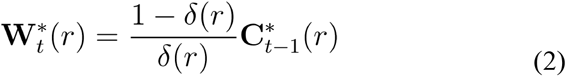

where *δ*(*r*) is a scalar ‘discount factor’. From this expression, it can be seen that *δ*(*r*) = 1 corresponds to a static model, while lower values allow dynamic connectivity estimates.

The prior hyperparameters 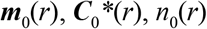, and *d*_0_(*r*) must be specified *a priori*. We used the same values as Costa et al. (2015b) which are 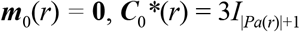 where *I* is the identity matrix, *n*_0_(*r*) = 0.001 and *d*_0_(*r*) = 0.001. We conducted prior sensitivity analyses and found these values are suitable for mean zero time series at each node and scaled (globally, over all time series and nodes) such that the average temporal standard deviation (SD) across nodes is 1 (it is crucial not to variance-normalize each node individually, as relative variance contains important directionality information).

The forecast distribution for *Y_t_*(*r*) given parents *Pa*(*r*) and past observations ***y***^*t*−1^ (*r*) follows a Student’s t-distribution (West and Harrison, 1997). This distribution allows us to evaluate the log likelihood of an observed time series by summing over time and regions. For an individual node *r* with parents *Pa*(*r*), the log likelihood, ie. the model evidence (ME), is

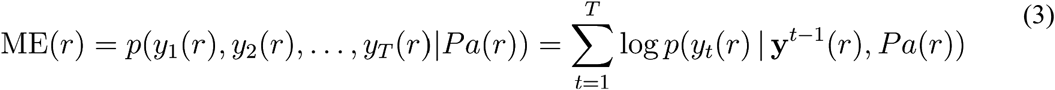

so that the model evidence for some model *m* with parents *Pa =* (*Pa*(1)*,…,Pa*(*n*)) is expressed as follows:

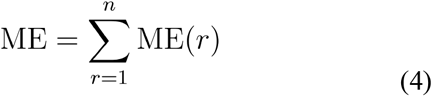

For each possible *Pa*(*r*), the ME is first maximised with respect to the discount factor *δ*(*r*) using a 1D grid search, *δ*(*r*) ∊ [0.5, 1]. While the MDM of Queen and Smith (1993) requires a DAG constraint, finding the highest ME with an acyclic configuration of parents (*Pa*(*1*) *,…, Pa*(*n*)), for DGM we simply find the maximum ME for each node individually (i.e. select the highest scoring set of parents). A model *m*_1_ is preferred to *m*_2_ if ME(*m*_1_) > ME(*m*_2_). Differences in ME correspond to log Bayes factors, and thus provide an interpretable magnitude of evidence for one model over another. In other words, a model is defined by the parents of a node; namely, whether there can be an influence or not. Given the model, it is possible to optimise the parameters in Equation 1, and evaluate the model evidence in Equation 3 and 4. The differences in model evidence can be used to directly compare models in terms of log Bayes factors (assuming that they have the same prior probability).

As DGM is fitting a collection of models, one per node, this collection may not necessarily correspond to a single statistical model (i.e. with a positive definite covariance posterior at every time point). This pseudolikelihood approach (Neville and Jensen, 2007) is an approximation to an ideal model, where (e.g.) latent nodes are introduced to account for reciprocal connections or other sources of cyclic behaviour. Each fitted DGM, thus, is best interpreted as a set of marginal node models that best describe the directed influence of parents on children nodes. This factorization of DGM model evidence by node is the same device used in regression DCM (Frässle et al., 2017).

### Example of a DGM with 3 nodes

Let’s assume a in a 3-node network with observed time series at the three nodes *Y_t_*(1), *Y_t_*(2), and *Y_t_*(3), see Figure 3. For example, node 3 has node 1 and node 2 as parents. Given the graphical structure, the model equations for the three nodes are written in Figure 3. The time-varying coefficients of the three edges e_1→2_, e_1→3_, e_2→3_ are 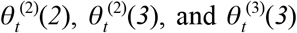, respectively. These estimates are for every time *t,* allowing the connectivity strengths to evolve over time.

### Model-comparison based pruning of reciprocal connections

We have considered an optional pruning process where bidirectional connections are converted to unidirectional connection as long as there is only modest reduction in model evidence. We find the optimal network, i.e. the configuration of parents (*Pa*(1),…,*Pa*(*n*)) that maximises the model evidence (ME). For each pair of nodes *i* and *j* with a bidirectional connection, we compare the model evidence for three possible models *m* (see also Figure 4):

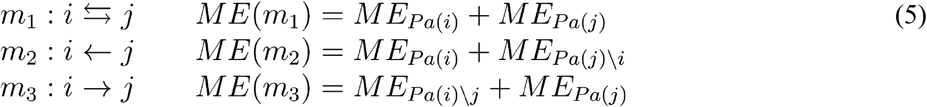

where *m*_1_ is the (ME-optimal) bidirectional edge, and the ME for these two nodes is the sum of the MEs of the two winning parent models node *i* and node *j*. *m*_1_ and *m*_2_ are the two models with unidirectional relationships, where each model evidence is likewise the sum of two MEs but with modified set of parents. For example in *m*_2_, where *i* is the child node, the likelihood (model evidence) is the sum of the ME of the winning parent model for *i* and the ME for *j* without *i* as a parent. Note that the total model evidence for the graph is a sum over all nodes, but since we will only change the model at these two nodes we can disregard the other *n - 2* contributions to the total ME for this comparison. The choice between *m*_1_, *m*_2_, and *m*_3_ is determined as follows: first, we define the evidence of the best unidirectional model as

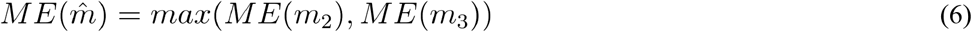

and then the model used is determined as

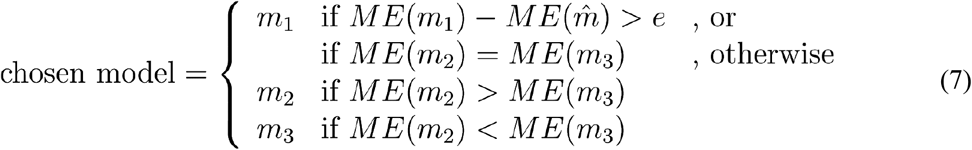

where *e* is a chosen penalty, interpreted as the log Bayes Factor comparing the original bidirectional model *m*_1_ to the best unidirectional model 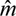. If the Bayes Factor for *m*_1_ is more than *e* we retain both the edges, otherwise, we chose a simpler uni-directed model. One can regard *e* as a prior over unidirectional versus bidirectional connections. In other words, if we use the value of *e* = 20, we are effectively saying that our prior odds for unidirectional connections over bidirectional connectivity is *exp*(20). For the simulation data, we used *e* = 20 for the 5-node network, and *e* = 10 for the 8-node network. For the 10 RSNs in human rs-fMRI, the estimated networks had strong evidence for unidirectional connections, with only 0.6% of edges altered with *e* = 20; as a result we used no pruning (*e* = 0). The same occurred with the mouse fMRI networks: we used no pruning as *e* = 20 altered only 7% of the edges.

**Figure 4.**
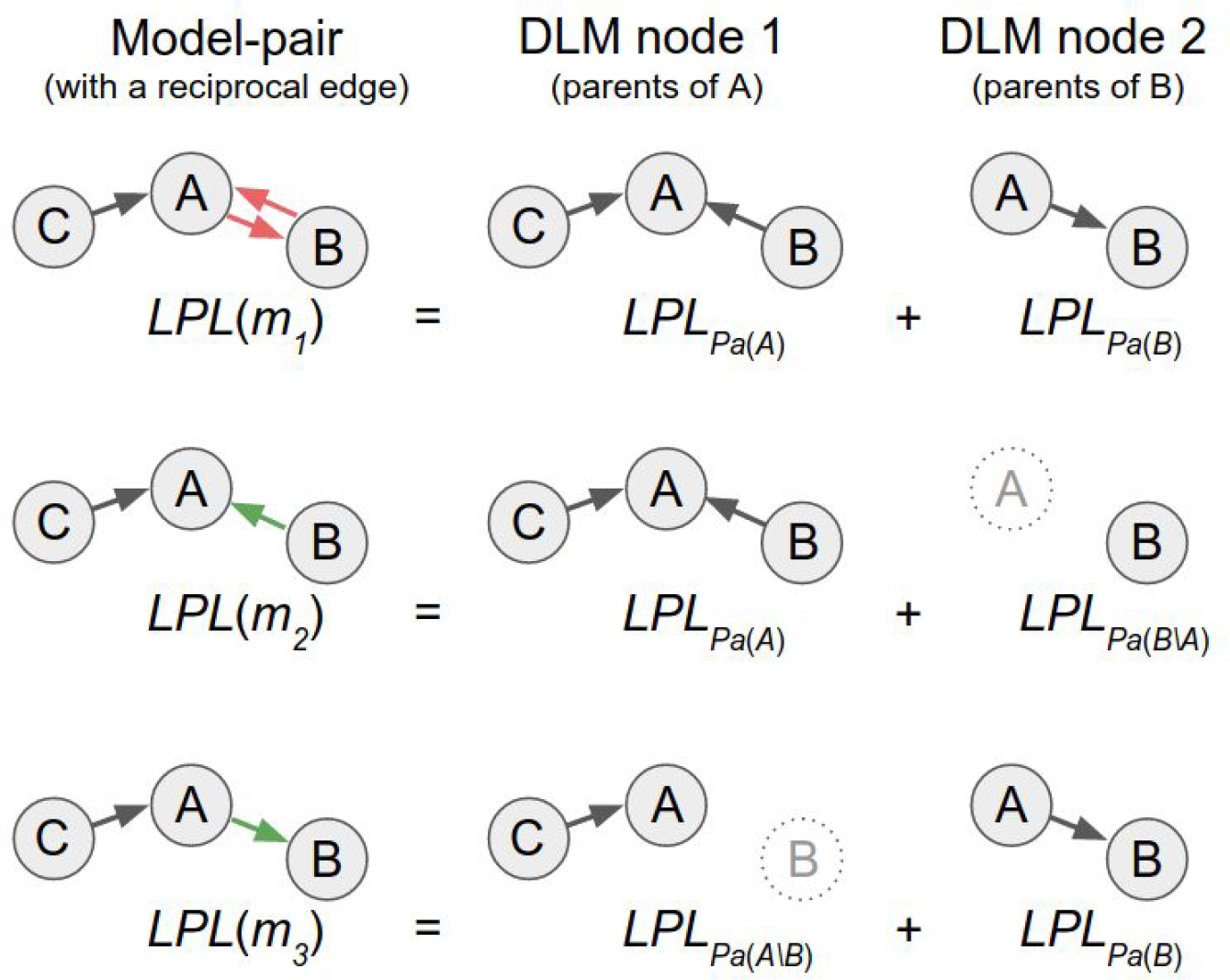
Pruning method’s model comparison approach, illustrated on a 3 node network. Each DGM with a bidirectional edge (red; here between node A and B) is a model-pair composed of two DLMs, one for node A (parents of A) and one for B (parents of B). We compare the model evidence (ME) between three models, the bidirectional (red) m_1_ and two unidirectional models (m_2_ and m_3_; green) where one or the other of the edges has been removed. We could choose a simpler model, and select the better unidirectional model if the difference in model evidence does not exceed an a priori defined value (in terms of a log Bayes factor).

### Statistical inference for edge consistency

After estimating the full networks for a dataset, we aim to test the reproducibility of the edges across simulated datasets or over real subjects and runs. We use a binomial test to identify edges that are more prevalent than expected by chance. First, we compute the sample proportion of edges at each directed edge from i to j

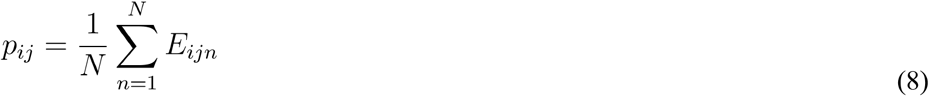

and define a null edge connection rate as

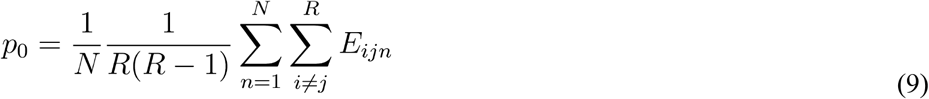

where E_ijn_ is a binary variable indicating the presence/absence of an edge for child node i and parent j from subject n, 1 ≤ i ≠ j ≤ R, n = 1,…,N. We then test each edge for edge prevalence that differs from the null edge connection rate, with null and alternative hypotheses:

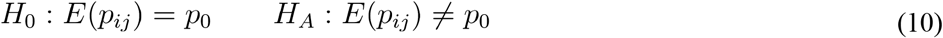

where *E(p_ij_)* is the true population probability of a specific edge with child *i* and parent *j* across *N* subjects. As we perform multiple testing across edges, we also adjust all *p*-values from the binomial test using false discovery rate (FDR) correction at the 5% level.

### Comparison with other directed methods

We compared DGM’s directionality estimates to a method proposed by Patel et al. (2006). Patel’s approach is based on pairwise conditional probability: An asymmetry in the probability of node A given node B, and B given A indicates a directed relationship. The model for the joint activation of each node pair is based on a multinomial likelihood with a Dirichlet prior distribution. As done in previous work (Smith et al, 2011), before calculating these conditional probabilities, the time series were scaled to an interval between 0 and 1 and then binarized with a cutoff at the 0.75 percentile. There are two measures derived from the conditional dependencies, the connection strength (κ) and the direction (τ). We implemented a permutation test for Patel’s κ that creates a distribution of κ values under the null hypothesis by shuffling the time series across subjects while keeping the nodes fixed. For each edge with a significant connection strength κ, we assigned the direction based on the sign of τ.

We also compared DGM to the pairwise Lingam method (Hyvärinen and Smith, 2013), an extension to the standard structural equation model that exploits non-Gaussianity of the data.

### Sensitivity and specificity

We investigated sensitivity and specificity for DGM and the various other network estimation methods to detect the true directed network in the simulations. For each simulation setting, we estimated networks from the simulated datasets and measured the number of true positives (TP), false positives (FP), true negatives (TN) and false negatives (FN), where sensitivity is the proportion true positives from condition positive [TP/(TP+FN)], and specificity is the proportion of true negatives from condition negative [TN/(TN+FP)]. Additionally, we computed “c-sensitivity” (Smith et al., 2011), the proportion of correct connections among all simulated datasets, regardless of the directionality.

### R package “DGM”

We released an R (RRID:SCR_001905) package “DGM” that implements DGM, Patel’s method and all the statistical procedures described above; the package is available on Github and on CRAN^3^. We used version 1.6.1 for all analyses. The number of parent models evaluated for each child node is 2^n-1^, and hence the total number of evaluations is n2^n-1^; for example a 25-node networks has 419 million possible models. Thus, we implemented some of the time critical functions in C++ using “Rcpp” (Eddelbuettel et al., 2011) and “RcppArmadillo” (Eddelbuettel and Sanderson, 2014). Currently, up to a 20-node network is computationally feasible (20 hours per node on a Intel Xeon CPU (E5-2630 v2) with 2.6 GHz), see Figure 1C. There is a future important extension that we will report separately: we have developed a greedy search algorithm that performs a stepwise search adding or removing a parent node in each step which drastically reduces the search space of all possible parent models (Harbord et al., 2016). The computations can be parallelized on the node level and example jobs are available for both Sun Grid Engine and Slurm on Github. “DGM” is free software, licensed under GPL3 and runs on all three major platforms (Linux, OS X and Windows).

## Results

### Network simulations

We evaluated the performance of the DGM to detect the true relationships^4^ of the non-stationary and stationary network nodes and compared to Patel’s and the Lingam method. On dynamic data the DGM (with *e* = 20) had higher sensitivity compared to Patel and Lingam (DGM: sensitivity and specificity, 70% and 79%; Patel: 42% and 82%; Lingam: 41% and 80%), see Figure 5A. DGM detected the 5 true edges in 14% of the cases, and 4 out of 5 in 56% of the time; Patel’s method detected 0% and 10%, and Lingam 0% and 8%, respectively. Looking at individual edges, the highest correct detection rate was 88% for 1→5 and the lowest was 52% for 4→5 for the DGM; Patel’s proportion of correct edges was lower, ranging from 24% (3→4) to 66% (1→5), and Lingam from 26% (1→2) to 60% (4→5), see Figure 5C. For DGM binomial significance tests on the edge proportions revealed all the 5 edges of the true underlying network for dynamic time series. Only the false positive edge 2→1 had a significant proportion of 62%, but was lower compared to the 78% for the true edge 1→2. With Patel’s method, only 3/5 of the true edges had proportions that reached significance, and there was a false positive edge at 1→3; Lingam discovered 3/5 edges but had 5 false positives and mainly estimated the inverse directions. When we increased the pruning to reach the same specificity as Patel’s method (82%) using *e* = 26, we found a sensitivity of 67% for DGM. Using no pruning (*e* = 0), DGM’s sensitivity was 84% and specificity was 64% (Figure 5A). For the DGM, the overall accuracy was best with *e* = 20 (76%) and *e* = 26 (78%), see Figure 5A. c-sensitivity was higher for DGM (89%) compared to Patel’s method (72%); we did not evaluate c-sensitivity for Lingam as this method only determined directionality. We also evaluated a longer simulation with DGM (60 min. instead of 10 min.) and found an increase in sensitivity to 91%, however, specificity was reduced to 60%; c-sensitivity was 98%.

**Figure 5.**
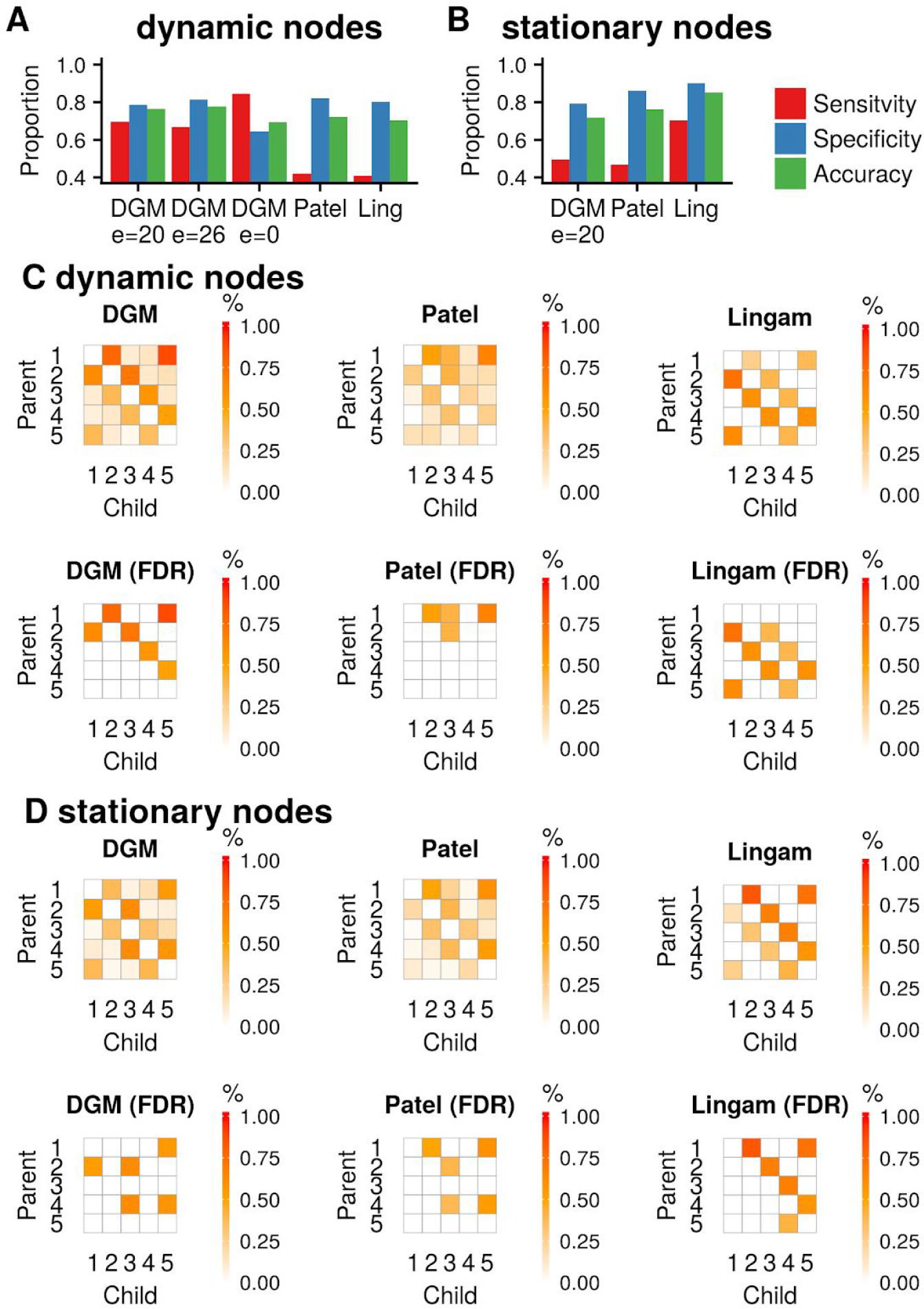
Sensitivity, specificity, and accuracy of the methods DGM, Patel and Lingam for dynamic **(A)** and stationary simulations **(B)**. **(C)** Proportions (top) and significant proportions (below, binomial test, 5% FDR threshold) for dynamic network simulations. **(D)** same as **(C)** for stationary network simulations.

**Figure 6.**
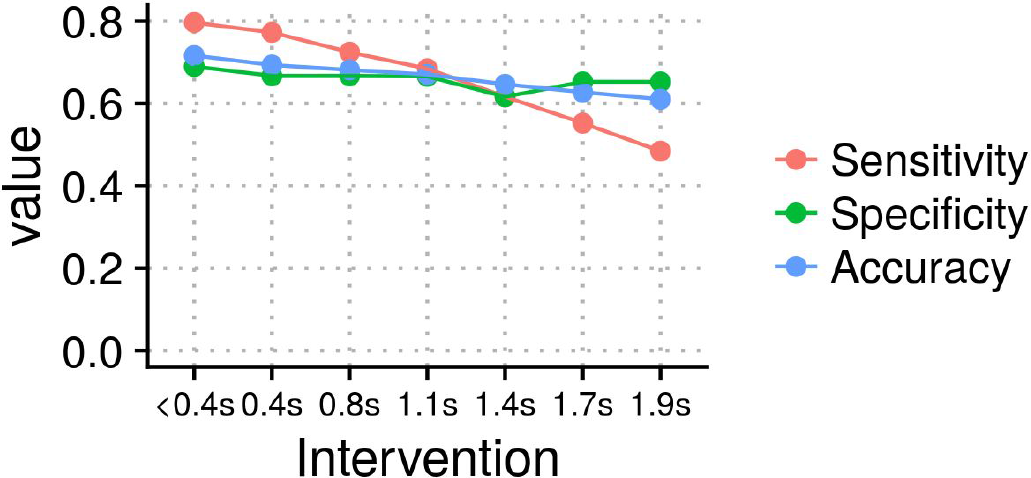
Sensitivity, specificity and accuracy for DGM (*e* = 20) for different node offsets (from 0.4 s up to 1.9 s total offset between the hemodynamic response of two nodes).

**Figure 7.**
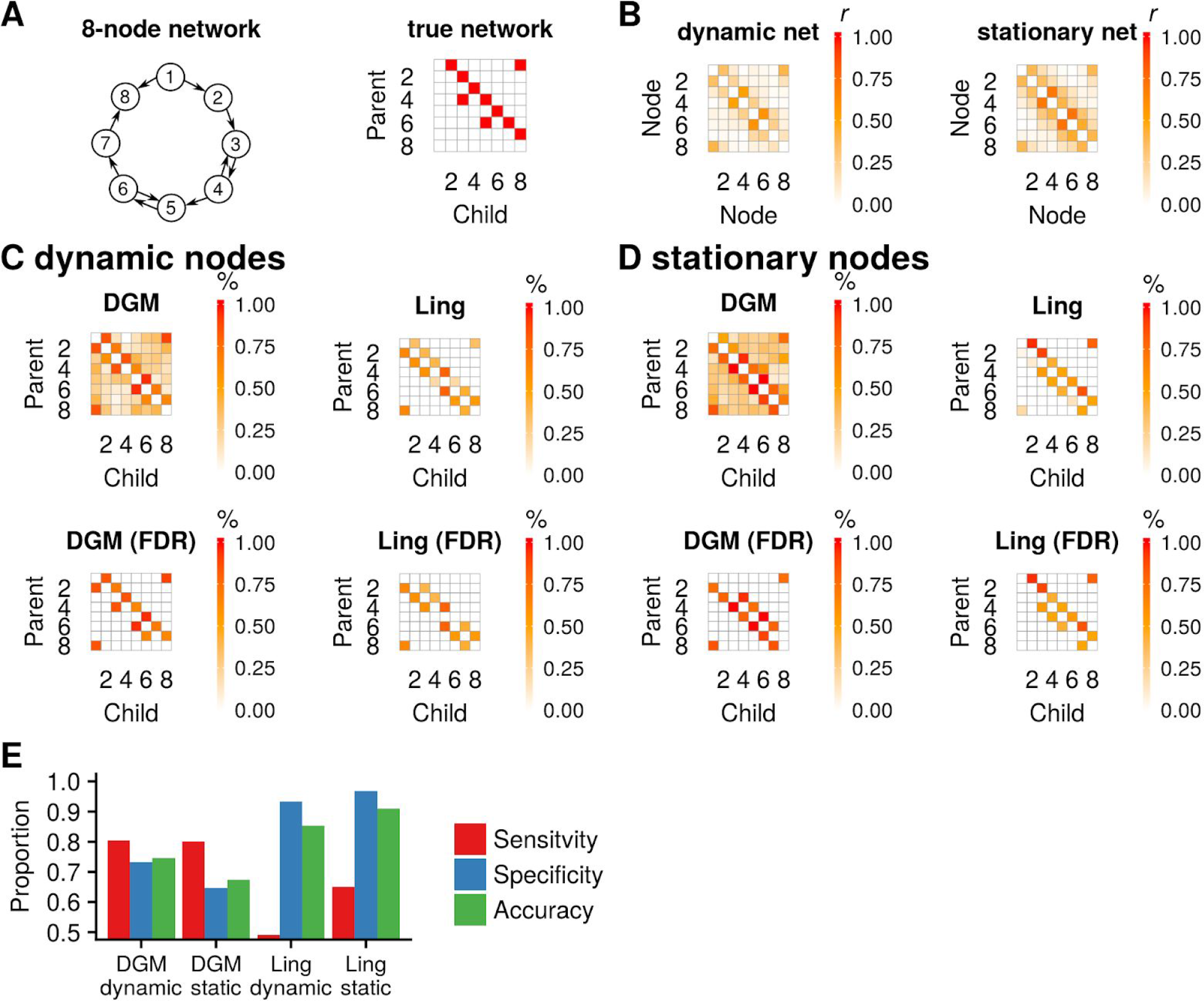
(A) The 8-node network with two reciprocal connections. **(B)** Pair-wise correlation matrix for both networks. **(C)** Proportions (top) and significant proportions (below, binomial test, 5% FDR threshold) for DGM and Lingam with dynamic network simulations. **(D)** same as **(C)** for stationary network simulations. **(E)** Sensitivity, specificity, and accuracy of the two network estimation methods for both types of simulations.

For stationary data, performance was best for Lingam with a sensitivity of 70% (DGM 50%; Patel: 47%), and a specificity of 90% (DGM: 79%; Patel: 86%), see Figure 5B. For DGM, the highest edge proportion was 62% for 2→3 and the lowest was 34% for 3→4, see Figure 5D. Using Patel’s method the proportion of correct edges ranged from 30% (3→4) to 60% (1→5), and for Lingam from 58% (4→5) to 84% (1→2). DGM discovered three of the five true edges that reached significant proportions with two significant edges that were false positive Patel showed 4/5 true edges that reached significance with one false positive edge, and Lingam recovered all true edges and one false positive.

For simulations with the systematic changes to the HRF delay (Figure 4), we found a sensitivity of 80%, 77% and 72% for the first three interventions, node offsets <0.4 s, 0.4 s, 0.8 s, respectively. Stronger interventions reduced sensitivity to 68%, 62% and 55%, for node offset 1.1 s, 1.4 s, 1.7 s, respectively. The strongest intervention with an offset of 1.9 s reduced sensitivity to 48%. Specificity ranged between 62%–69%.

We looked at the discount factors *δ*(r), the temporal smoothness of the connectivity strengths ***θ****(r)* for nodes with at least one parent. Median *δ(r)* across the 5 nodes was 0.86 (0.84–0.89) (Q1–Q3) for the dynamic data Sim22, and 0.67 (0.66–0.67) for the dynamic simulations with the least intervention (< 0.4 s) and 0.71 (0.65–0.75) for the one with the strongest intervention (1.9 s). For the stationary simulation (Sim1), a median *δ(r)* of 0.994 (0.992–0.996) was fitted.

We also tested a 8-node network with reciprocal connections (Figure 7A). The reciprocal connections showed higher correlation compared to the unidirectional edges, which was expected (Figure 7B). On dynamic data the DGM (with e = 10) had higher sensitivity compared to Lingam (DGM: sensitivity and specificity, 81% and 73%; Lingam: 49% and 93%), see Figure 5E. However, note here that Lingam cannot detect a relationship between two nodes, only the direction of the edge, and therefore we informed Lingam with the edges that have a true connection. For DGM binomial significance tests on the edge proportions revealed all the 10 edges of the true underlying network for dynamic time series with 3 false positives. With Lingam, only 7/10 of the true edges had proportions that reached significance, and there were 5 false positive edges. On stationary data the DGM (with *e* = 10) had higher sensitivity compared to Lingam (DGM: sensitivity and specificity, 80% and 65%; Lingam: 65% and 97%), see Figure 5E. After significance testing, DGM revealed all the 5/10 edges with 6 false positives, and Lingam recovered all the 10 edges that are true with 2 false positives, see Figure 7D.

### Human resting-state fMRI

The ICA-based resting-state networks from the HCP data had consistent edges in in up to 60% of the participants. The cerebellar, auditory, and also the visual medial networks were child nodes to other RSN’s (Figure 8). The visual medial network receives input most prominently from the two other visual networks (visual pole and visual lateral) and the DMN. The visual lateral network is reciprocally connected with the visual medial network and also receives input from the occipital pole. Notably, the estimated networks look very similar across runs. Figure 9 shows the within-subject consistency of the estimated networks, both in terms of presence (Figure 9A) and absence (Figure 9B) of edges. We quantified the edges that occurred in three out of the four runs, and those that occurred in all of the runs. We found that some of the edges described above are reproducible in up 50% of the participants in three runs, and in 30% of the participants in all the four runs. This applied to the reciprocal visual medial and lateral connections and the cerebellum inputs from the DMN, the visual medial, and the auditory network. In terms of reliably absent edges, the visual occipital pole, the DMN, both the right and left frontoparietal network, and also partly the executive control network had consistently no other RSNs as their parents (Figure 9B).

**Figure 8.**
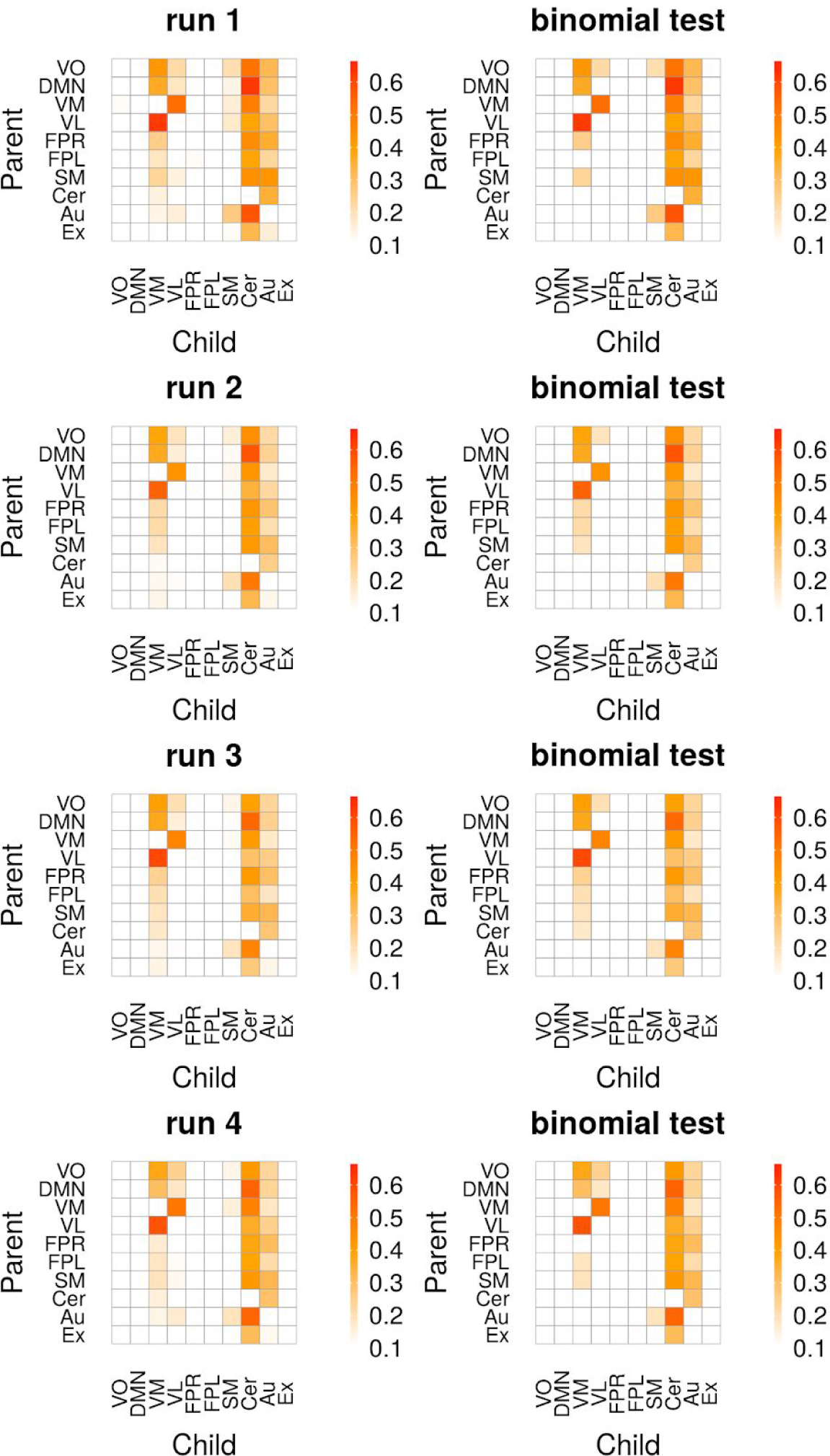
Proportions of edges in participants (*N* = 500) from 4 runs (top to bottom) during resting-state of approximately 15 minutes. Left shows all proportions, right shows proportions for significant edges (binomial test, 5% FDR threshold). Legend: VO, visual occipital pole; DMN, default mode; VM, visual medial; VL, visual lateral/ventral; FPR, frontoparietal right; FPL, frontoparietal left; SM, sensorimotor; Cer, cerebellum; Au, auditory; Ex, executive control (including thalamus).

**Figure 9.**
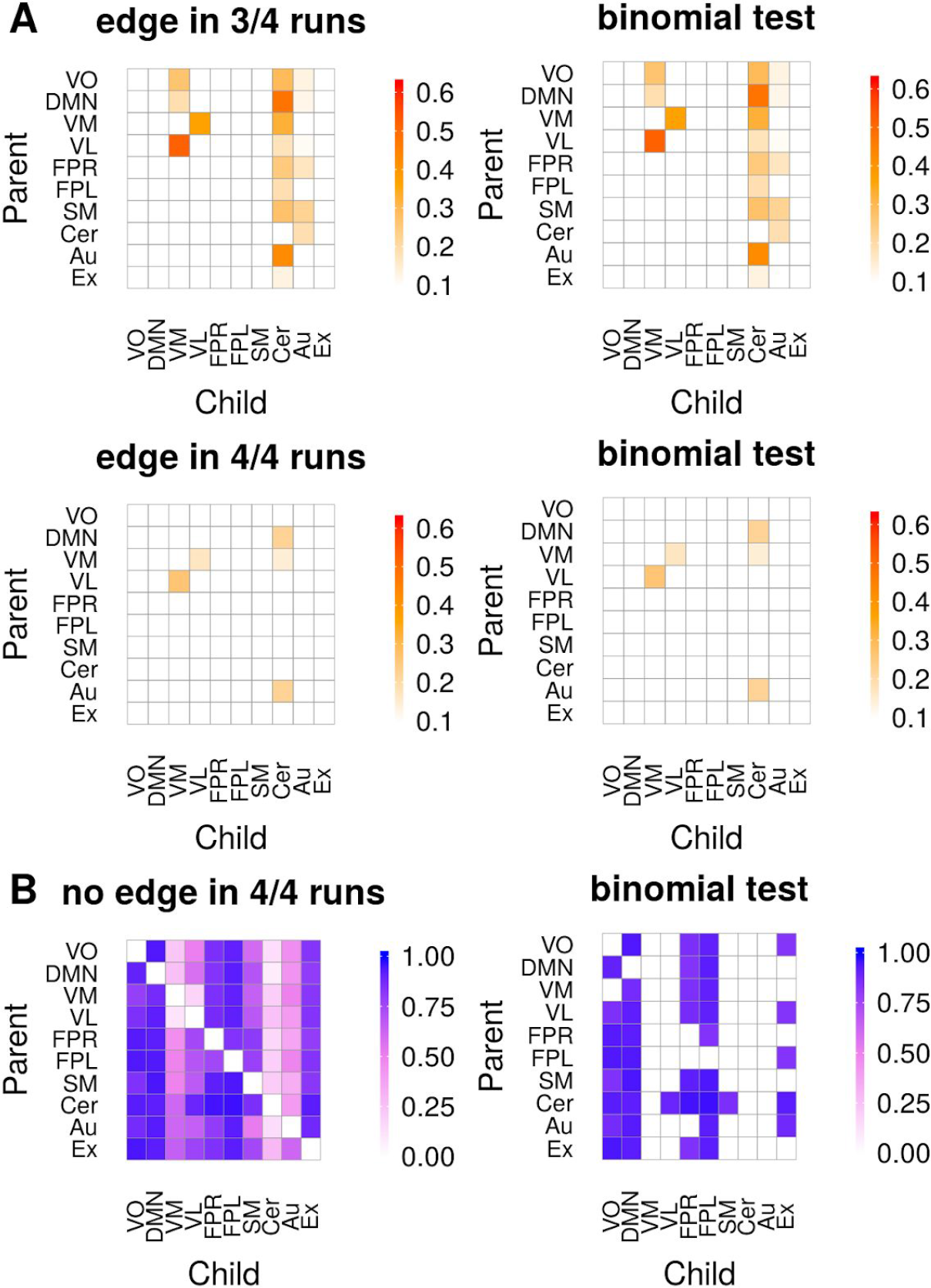
**(A)** Proportions of edges (red) from 500 participants that reoccur in three of the four runs, or in all the four runs (within-subject reproducibility). **(B)** Proportion of participants that had no edge (blue) across all runs. Left shows all proportions, right shows proportions only for significant edges (binomial test, 5% FDR threshold).

The discount factors *δ*(r) of the connectivity strengths for nodes with at least one parent had a median (Q1-Q3) across the 10 RSNs of 0.80 (0.74–0.88) in run 1; the runs 2-4 showed similar values.

### Mouse resting-state fMRI

We used DGM to investigated dynamic directed functional networks in rs-fMRI data from 16 mice under light anesthesia. We analyzed time series from two networks: the first network has 8 nodes which are regions that are functionally connected with the dentate gyrus of the hippocampus (DG). Directed anatomical connectivity based on tracer data suggests that hippocampal areas (DG and CA1) receive projections from the entorhinal and other associative cortices (Weilbächer and Gluth, 2016), the nucleus accumbens and the amygdala, and project to prefrontal areas (Eichenbaum, 2017; Hintiryan et al., 2016; Jin and Maren, 2015; Oh et al., 2014). We found that the DG received received inputs from several nodes, but missed significance. However, a large proportion, 94%, of mice showed directed connectivity from the orbital cortex (OrbM) to the cingulate cortex, see Figure 11A. We also found multiple parent regions of the OrbM area, including the DG (63% of mice). Furthermore, the node CA1 (63% of the mice) and DG (75%) were parent nodes of the cingulate cortex.

**Figure 10.**
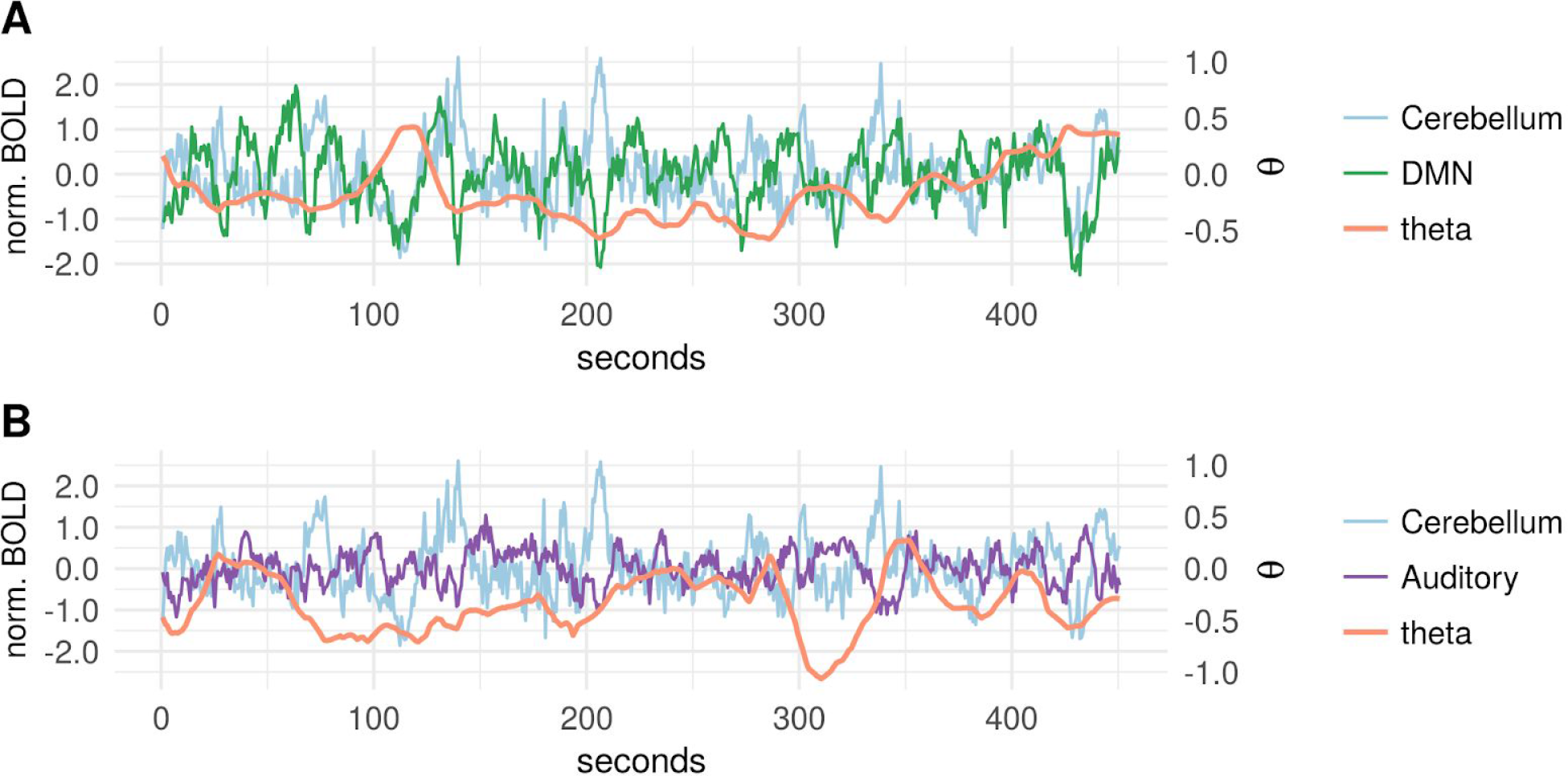
Example of time-varying connectivity estimates ***θ****_t_*(*r*) for two parents of the cerebellar network, subject 108525, T=0–450 s. **(A)** Cerebellar network time series (blue), plotted with the default mode network (green) and its directed influence reflected by the ***θ****_t_*(*r*) time series (orange); note the transient positive dependence around 70 seconds contrasted with otherwise negative dependence. **(B)** Cerebellar network time series plotted with the auditory network (violet) and its ***θ****_t_*(*r*) connectivity; the influence of the auditory network is generally negative.

**Figure 11.**
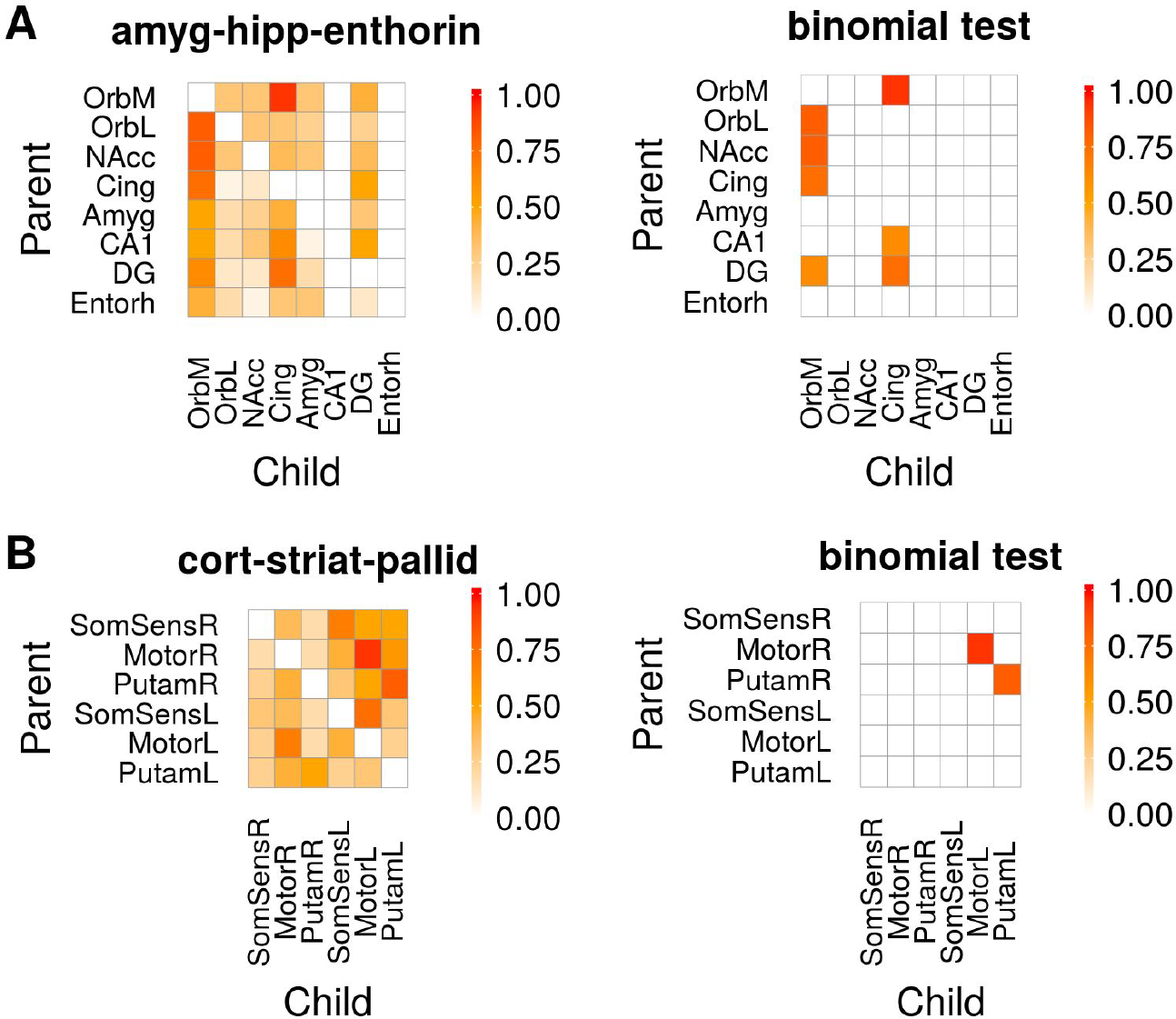
Proportion of edges in two functional networks of mice (*N* = 16). **(A)** amygdala-hippocampal-entorhinal network and **(B)** an cortico-striatal network. Left shows all proportions, right shows significant proportions of edges (binomial test, 5% FDR threshold). Legend: orbital area, medial (OrbM) and lateral part (OrbL); nucleus accumbens (NAcc); anterior cingulate area, dorsal part (Cing); lateral amygdalar nucleus (Amyg); cornu ammonis area 1 (CA1); dentate gyrus (DG); entorhinal area, lateral part (Entorh); primary somatosensory area, barrel field right and left (SomSensR, SomSensL); primary motor area right and left (MotorR, MotorL); caudoputamen right and left (PutamR, PutamL).

The second network contains six nodes that communicate with the motor and somatosensory barrel field areas. Based on axonal/white-matter connections, the cortical motor and somatosensory areas project to the striatum, and also contralateral flow between the areas can be expected (Lee et al., 2016). We found the highest consistency across mice in the contralateral connectivity from the right to the left motor cortex (94%), and from the right to the left putamen (81%), see Figure 11B.

The discount factors *δ*(r) of the connectivity strengths for nodes with at least one parent had a median (range) across nodes of 0.99 (0.990–0.994) in the 8-node network, and 0.99 (0.955–0.995) in the 6-node network.

## Discussion

Dynamic graphical models (DGM) are a collection of dynamic linear models (DLM) at each node where we treat parent nodes as covariates in a dynamic multiple regression. Even though the (graphical) model structure is static once we optimized the family of parental nodes, we allow for time-varying correlations in the observed fMRI time-series. We extended previous work by (Costa et al., 2015a), relaxing the DAG constraint and allowing for cycles. Furthermore, we introduced systematic lags to test robustness against hemodynamic confounds and we could successfully invert time-varying node relationships in simulation data generated by a the Smith et al. (2011) generative model based on a deterministic DCM. In determining directed relationships we were partly successfully, but this remains a challenging task. We could establish consistent relationships across a large number of subjects that demonstrated top-town influence from the DMN to the cerebellum and limbic regions, as well as to temporal and memory related areas; and we found a reciprocal connection of the visual medial and visual lateral areas. In the future, DGM has the potential to estimate large networks with 100 of nodes (by optimization of network discovery) which can be suitable for further analyses with graph theory (Bullmore and Sporns, 2009; Fornito et al., 2016). Hopefully, this device allows to study pathological network topologies of brain disorders in the future (Fornito et al., 2015), but further validation of the methods are required.

We found that DGM had a sensitivity and specificity between 70%–80% for the 5 and 8-node network with nonstationary node relationships. Patel’s method suffered low sensitivity to detect a relationship between nodes but can establish the correct directionality in a majority of edges. Lingam oriented the network correctly, but only in the stationary case. However, one limitation here is that Lingam in not a full model, i.e. the approach requires to know the edges a priori and then orients those based on non-gaussian properties.

DGM showed lower sensitivity with stationary data in the 5-node network but not for the 8-node network. This is interesting as DGM has the ability to fit a fully stationary model, and indeed such a stationary model was generally fitted with the stationary data. However, the directionality structure then often suffers in this case and it appears the dynamic information in the data is highly relevant in the determination of directionality.

The main finding for the human resting-state data is that the cerebellar network is most consistently receiving input from many other networks. A meta-analysis demonstrated that the cerebellum is not only relevant in fine motor skills but also involved in cognitive functions, such as working memory, language, emotions, and executive control (Stoodley and Schmahmann, 2009), and subregions of the cerebellum have been functionally related to RSNs (Habas et al., 2009). It is important to note that the cerebellar component we used also included appreciable weights for the thalamus and the limbic system, most prominently the amygdala, hippocampus, and the anterior cingulate gyrus. These regions received input from the DMN, a finding which was consistent across subjects and also within-subject across the four runs. This could reflect a intrinsic dynamic coupling of regions associated with memory and emotion with the DMN (Sestieri et al., 2011). The auditory component not only involved the superior and middle temporal gyrus but also language related left frontal structures and the temporal poles which are highly relevant in semantic memory (Galton et al., 2001). We found that the auditory/language network also receives input from various other networks which could reflect the dynamic access to the language and semantic networks. Furthermore, we found that visual networks have high consistencies in dynamic coupling, such as between the visual medial and the visual lateral network.

We used DGM to explore directed functional connectivity in the mouse brain. We found some highly reproducible edges (in up to 94% of the mice), and could successfully determine that CA1 and DG are feeding to the cingulate cortex; this finding is consistent with studies that use viral tracers to determine the directed anatomical connectivity (Hintiryan et al., 2016; Oh et al., 2014). A limitation here is that we compare functional connectivity with findings from anatomical connectivity which are conceptually different and certainly do not have a one-to-one relationship, however, there is evidence from functional studies that the hippocampus and the anterior cingulate cortex are prominent areas of the limbic circuit and the DMN.

In DGM we use a discount factor δ (Petris et al., 2009b; West and Harrison, 1997) that sets the temporal smoothness of the connectivity strength *θ* for each node; this can range from a maximally smooth path (i.e. a constant *δ*(*r*) = 1) to a highly variable path (*δ*(*r*) = 0.5). For each node, and for each set of parents, we find the DF that optimizes the model evidence (ME), and that optimised ME is used to identify the optimal set of parents. We found that the δ was higher in Sim22 compared our own dynamic simulations. A possible explanation is that we used a higher sampling rate (lower TR = 2) which preserves more dynamic information in the time series. In the mouse data, we found a δ near 1 suggesting that DGM were not driven by dynamic relationships and was mostly stationary. This may be related to the fact that the mice have been anesthetized (Barttfeld et al., 2015), however, other studies were successful to determine dynamic relationships in anesthetized animals (Hutchison et al., 2013b).

Various methods exist to establish directed relationships: Granger causality for fMRI, as a lag-based method, can be confounded by the variability in the HR of different brain areas. Latency correction can improve detectability (Wen et al., 2013), but the magnitude of this confound is usually unknown in real data. DCM (Friston, 2009; Friston et al., 2003) implements a biophysical model and can, in principle, infer upon the latent neural mechanisms, but has a high computational effort. However, recent developments allow DCM to study larger networks and also resting-state fMRI (Frässle et al., 2017; Razi et al., 2017). In future work, we would like to establish construct validity between DGM and spectral DCM (spDCM) for resting state (Friston et al., 2014a; Razi et al., 2015). In the current paper, we focused on comparing DGM to Patel’s method and Lingam within the scope of purely observational models of directed functional connectivity. In contrast, spDCM, which aims to infer effective connectivity, has a different agenda which is to infer causal influences between different neuronal populations. spDCM is a hierarchical model with two sets of neural and observation (hemodynamic) equations. spDCM assumes neural fluctuations to have a power law (scale-free) distribution whereas the deterministic (forward) DCM for our simulations had an exogenous input (i.e., no endogenous fluctuations). Therefore, our current simulations are not compatible with the generative model employed in spDCM. Another method to determine directed functional connectivity is Group Iterative Multiple Model Estimation (GIMME), a structural equation model (Gates and Molenaar, 2012). Here, information on an estimated group-level network is used to improve network structure estimates on the subject-level. The method showed a high sensitivity and accuracy (both above 80%), but we have not evaluated it here as it does not provide subject-level estimates.

Considering the overall accuracy as the only measure of evaluation can be biased, especially with a sparse network with only 5 out of 20 possible edges. With Patel’s method for example, even though detection of edges (sensitivity) was poor, overall accuracy was high due to the high true negative rate. We evaluated sensitivity for both (1) the successful detection of a relationship between nodes and (2) the correct estimation of the directionality simultaneously; while other work (Costa et al., 2015a; Smith et al., 2011) looked separately at (1) and (2) and found good directionality estimates (0.60–0.70; d-accuracy) but only under the assumption of (1) being successful. We specifically addressed this issue with a randomization test for Patel’s connection strength κ and then assigned the directionality based on the sign of τ.

We stress that the DGM is a heuristic approach, and demands thorough simulations and critical evaluations in the context that it is to be applied, as we have done here with resting fMRI. A particular problem to fMRI is the variation of the hemodynamic response across brain regions that may a confounding factor for lag based methods to estimate directionality. DGM, however, is driven by instantaneous relationships between nodes but does consider some information from the past to regularise state variables (Equation 1; system equation). We created network simulations with systematic changes on the balloon model priors to increase the lag at parent nodes and decrease the lag at child nodes. We found that if the total offset between a node pair is in the problematic direction (i.e., which would cause Granger to infer incorrect directionality) but below one second, the DGM’ sensitivity remains above 70%. If this offset is further increased, sensitivity is declining, but specificity remains intact. Thus, such edges cannot be detected anymore, but incorrectly reversed directionality between the node-pair is not inferred (i.e. the false positive rate).

Even though the main focus of this paper is on data from fMRI, DGM are not restricted to this type of data and can potentially be applied in electroencephalography or magnetoencephalography that has a much higher temporal resolution (Baker et al., 2014).

## Conclusions

Our results demonstrate that DGM can discover directed functional connectivity that constitutes reproducible edges in human and mouse rs-fMRI. We observed that in humans the cerebellar/limbic network consistently receives information from other networks. In network simulations, DGM demonstrated a sensitivity of 72%–77% for dynamic time-series, even in the presence of systemic hemodynamic lag confounds. DGM is a novel and promising approach that provides news insights into directed dynamic relationships in functional connectivity.

## Software and Data Availability

The source code of the R package “DGM” is available at https://github.com/schw4b/DGM. Data and analyses can be found at https://github.com/schw4b/DGM-Sim.

## Acknowledgements

We thank Karl Friston and Adeel Razi for their expert advice and an in-depth discussion of this work. Data were provided in part by the Human Connectome Project, WU-Minn Consortium (Principal Investigators: David Van Essen and Kamil Ugurbil; 1U54MH091657) funded by the 16 NIH Institutes and Centers that support the NIH Blueprint for Neuroscience Research; and by the McDonnell Center for Systems Neuroscience at Washington University.

## Footnotes

1 Number of directed graphs with no self-loops, https://oeis.org/A053763.

2 http://www.fmrib.ox.ac.uk/datasets/netsim/

3 https://github.com/schw4b/DGM

4 Although we refer to the known connections generating the data as ’true’ we do not necessarily imply that these are the ’best’ explanations of the observed data. It is commonplace to generate data for which there are simpler explanations. In other words, it is possible that our analyses identified more parsimonious explanations for particular realisations of the data.

