## Supplementary Materials for "Directed functional connectivity using dynamic graphical models"

#### Supplementary Table S1

**Supplementary Table S1:** Description of the ten resting state networks used.

| Component from HCP parcellation time series from NetMats d25 | Corresponding RSN from Smith et al. (2009) | Spatial correlation (Pearson's r) | Description (Harvard-Oxford and Cerebellar Atlas) | Label of network (Abbreviation) |
|---|---|---|---|---|
| 1 | 2 | .57 | Occipital pole | Visual occipital (VisOcc) |
| 2 | 4 | .57 | precuneus, cingulate g., superior frontal g., middle frontal g., middle temporal g., temporal pole, lateral occipital cortex (c.) hippocampus, frontal orbital c., cerebellum (crus I, II) | Default mode (DMN) |
| 3 | 1 | .63 | Lingual gyrus, cuneus, cerebellum (VI), occipital pole, precentral g. | Visual medial (VisMed) |
| 4 | 3 | .55 | lateral occipital c., temporal occipital fusiform c. | Visual lateral (VisLat) |
| 5 | 9 | .57 | Various areas in frontal and parietal c. right | Frontoparietal right (FrontParR) |
| 10 | 10 | .53 | Frontal and parietal areas left | Frontoparietal left (FrontParL) |
| 14 | 6 | .39 | Precentral g., postcentral g., operculum, cerebellum (V, VI, VIII) | Sensorimotor (SensMot) |
| 18 | 5 | .36 | Cerebellum (posterior lobe), amygdala, thalamus, hippocampus, cingulate g. (anterior) | Cerebellar (Cerebell) |
| 19 | 7 | .41 | Superior temporal g., operculum, supramarginal g., thalamus, superior frontal g., hippocampus, temporal pole | Auditory (Aud) |
| 21 | 8 | .42 | medial frontal areas, anterior cingulate, paracingulate | Executive control (Exec) |

### Supplementary Table S2

**Supplementary Table S2**: Mean (SD) peak of HRF after stimulus onset in seconds for different interventions and nodes across 50 simulated datasets.

| Simulation name (total offset between nodes) | $\tau$ node 1 and 4 | $\tau$ node 2 and 5 |
|---|---|---|
| < 0.4 s | 0.98 | 0.98 |
| 0.4 s | 1.27 | 0.75 |
| 0.8 s | 1.57 | 0.61 |
| 1.1 s | 1.96 | 0.49 |
| 1.4 s | 2.35 | 0.41 |
| 1.7 s | 2.74 | 0.35 |
| 1.9 s | 3.14 | 0.31 |

### Supplementary Table S3

**Supplementary Table S3**: Mean (SD) peak of HRF after stimulus onset in seconds for different interventions and nodes across 50 simulated datasets.

| | < 0.4 s | 0.4 s | 0.8 s | 1.1 s | 1.4 s | 1.7 s | 1.9 s |
|---|---|---|---|---|---|---|---|
| Node 1 | 4.07(±0.25) | 4.30(±0.31) | 4.53(±0.32) | 4.68(±0.31) | 4.90(±0.32) | 5.15(±0.40) | 5.36(±0.45) |
| Node 2 | 4.10(±0.29) | 3.89(±0.31) | 3.76(±0.33) | 3.67(±0.25) | 3.61(±0.25) | 3.57(±0.26) | 3.55(±0.25) |
| Node 3 | 4.18(±0.36) | 4.19(±0.31) | 4.19(±0.31) | 4.19(±0.31) | 4.19(±0.31) | 4.19(±0.31) | 4.19(±0.31) |
| Node 4 | 4.28(±0.30) | 4.47(±0.36) | 4.78(±0.35) | 5.03(±0.31) | 5.13(±0.29) | 5.27(±0.38) | 5.46(±0.42) |
| Node 5 | 4.05(±0.35) | 3.86(±0.25) | 3.71(±0.25) | 3.57(±0.28) | 3.47(±0.25) | 3.43(±0.23) | 3.42(±0.23) |

### Supplementary Table S4

**Supplementary Table S4**: Peak offsets in seconds relative to first simulation (< 0.4 s)

| | 0.4 s | 0.8 s | 1.1 s | 1.4 s | 1.7 s | 1.9 s |
|---|---|---|---|---|---|---|
| Node 1 | +0.23 | +0.46 | +0.62 | +0.84 | +1.09 | +1.29 |
| Node 2 | -0.21 | -0.34 | -0.43 | -0.49 | -0.53 | -0.55 |
| Node 3 | +0.01 | +0.01 | +0.01 | +0.01 | +0.01 | +0.01 |
| Node 4 | +0.19 | +0.50 | +0.75 | +0.85 | +0.99 | +1.18 |
| Node 5 | -0.19 | -0.35 | -0.49 | -0.58 | -0.63 | -0.63 |

### Supplementary Table S5

**Supplementary Table S5**: Total offset in seconds between nodes with increased/decreased lags.

| | 0.4 s | 0.8 s | 1.1 s | 1.4 s | 1.7 s | 1.9 s |
|---|---|---|---|---|---|---|
| node 1 and 2 | 0.44 | 0.80 | 1.04 | 1.32 | 1.62 | 1.85 |
| node 4 and 5 | 0.38 | 0.84 | 1.24 | 1.44 | 1.62 | 1.82 |
| node 1 and 5 | 0.43 | 0.81 | 1.10 | 1.42 | 1.71 | 1.93 |
